# Axenic Aedes *aegypti* develop without live bacteria, but exhibit delayed development and reduced oviposition

**DOI:** 10.1101/264978

**Authors:** Maria A. Correa, Doug E Brackney, Blaire Steven

## Abstract

The mosquito gut microbiome plays an important role in mosquito development and fitness, providing a promising avenue for novel mosquito control strategies. Here we present a method for rearing axenic (bacteria free) *Aedes aegypti* mosquitoes, which will greatly facilitate mechanistic studies documenting the structure and function of the microbiome. Through feeding sterilized larvae agar plugs containing attenuated *Escherichia coli*, mosquito development was observed in the absence of living bacteria. Axenic larvae were capable of full development into adults, which laid eggs that were subsequently hatched. However, axenic mosquitoes exhibited delayed development time and reduced egg clutch size in comparison to bacterially colonized mosquitoes. These findings suggest that mosquito development is not dependent on live bacteria, but their phenotype is modulated by the presence of microorganisms. This axenic system offers a new tool in which the mosquito microbiome can be systematically manipulated for a deeper understanding of microbiome host interactions.

## Introduction

It is increasingly clear that most, if not all, multicellular organisms live in association with a complex assemblage of microorganisms (i.e. microbiome) composed of bacteria, viruses, fungi, and archaea. While these communities can be found in every habitable organ, for most complex organisms the vast majority of microbes reside in the digestive tract. Because of the biomass and complexity of the indigenous gut microbiome, as well as its close association with the host, it is frequently considered an additional major organ^1,2^. Consequently, the influence of microbiota on host biology has garnered considerable attention^3,4^. These studies have revealed a link between the microbiome and a wide array of disease states in mammals, including obesity^5^, diabetes^6^, and autism^7,8^. Furthermore, the microbiome has been implicated in playing a significant role in the development and function of the immune system and autoimmune disorders^9,10^.

Invertebrates also harbor a diverse microbiome^11,12^ that has been linked to a number of phenotypic outcomes, such as host-mating preference^13^ and embryonic development^14^. It is clear from these studies that the microbiome can have profound effects on host physiology and health. Mosquitoes are important disease vectors for a number of human pathogens that include arboviruses, protozoa, and nematodes that pose a significant public health threat. Due to the lack of an effective vaccine for many of these pathogens and an increase in insecticide resistance in mosquitoes, the development and implementation of novel mosquito control strategies will be necessary to curtail their public health impact. The mosquito microbiome is emerging as a potential tool in this effort^15^. A number of descriptive studies, primarily focused on the bacterial components of the microbiome, have determined that it is relatively simple, typically composed of 10-70 bacterial strains, the majority of which are members of the phylum Proteobacteria,specifically the family *Enterobacteriaceae^16,17^.* Based on the similarity in composition between the microbiome of mosquito larvae and the water they inhabit, it has been proposed that mosquitoes largely acquire their gut microbiota from the aquatic environment^18^. Further evidence suggests that at least some of the larval microbiome is transstadially transmitted to the adult after pupation^19^. While these studies demonstrate that environmental microorganisms readily colonize mosquitoes and these associations can be stable over the entire lifespan, the role these microbes play in mosquito development and biology is less clear.

Most mosquito-borne pathogens must infect or pass through the mosquito midgut prior to being transmitted. Not surprisingly, the complex interplay between pathogens, the mosquito midgut, and its associated microbiome have garnered considerable attention. For instance, it is known that bacterial load and/or microbiome community composition can significantly affect *Anopheles spp.* mosquito susceptibility to *Plasmodium* infection^20,21^ and *Aedes aegypti* susceptibility to dengue virus is influenced by the intestinal microflora^22,23^. Microbiota have differing effects on vector competence in mosquitoes, with particular isolates either positively or negatively influencing mosquito infection rates depending on the species or bacterial strain^22,24-26^. Taken together, these observations demonstrate that the composition and structure of the microbiome can affect the ability of mosquitoes to acquire and transmit disease. Yet, because no current method exists to systematically manipulate the microbiome these studies are by definition correlational in nature. Furthermore, it is difficult to determine the impact of the microbiota on mosquito-pathogen interactions because many of these studies have relied upon antibiotic “clearance” of the bacterial communities. Recent reports show that the mosquito microbiome often contains antibiotic resistant bacteria^27^, antibiotics do not fully clear the gut microbiota, but rather cause a dysbiosis^28^, and extended use of antibiotics can cause toxicity and mitochondrial dysfunction^29^. Consequently, these models cannot be truly considered “bacteria free” and do not address possible interactive effects between bacterial reduction and antibiotic exposure on mosquito biology and vector competence.

Systematic manipulation of the mosquito microbiome would be greatly facilitated by the existence of a microbiome-free, or axenic, mosquito^30^. Furthermore, the development of an axenic model could act as a blank template on which a microbiome of known composition could be imprinted, also known as a *gnotobiotic* organism^31,32^. Axenic rearing techniques have already been developed for a number of model organisms, including *Drosophila melanogaster*, *Caenorhabditis elegans,* and mice^30^. Early attempts to rear axenic mosquitoes reportedly obtained adults free from bacteria using a growth media of essential vitamins and nutrients^33^. However, these studies lacked the modern molecular based techniques that can detect microorganisms that are recalcitrant to laboratory cultivation; as is frequently cited, the majority of microorganisms in nature are unculturable^34^. In this regard, there is some uncertainty as to whether these mosquitoes were truly axenic. In fact, a series of recent studies have reported that mosquitoes require a live bacterial symbiont for development^27,35,36^. Yet the studies describing the necessity of live bacteria generally ignored the role of microflora in supplying essential nutrients to the host. Thus, there is somewhat of a contradiction in the literature; either mosquitoes are unique from *Drosophila* and *C. elegans* in requiring a live bacterial symbiont for development, or nutritional conditions sufficient to rear axenic mosquitoes have yet to be documented.

In this study, we tested a variety of reported methods and developed novel practices for the rearing of *Aedes aegypti* mosquitoes free of living bacteria. A means to rear axenic mosquitoes was achieved by hatching larvae from surface sterilized eggs fed on agar plugs containing sonicated and heat-inactivated bacteria. Biometric comparisons revealed that developmental times differed between axenic mosquitoes and gnotobiotic mosquitoes colonized by *Escherichia coli.* Axenic mosquitoes also had a significant reduction in egg clutch size compared to their microbiome-colonized cohorts. The data presented here represents a methodological advancement in the field of mosquito microbiome research and provides a much-needed tool to elucidate the role of microbiota in mosquito physiology and pathogen susceptibility. Furthermore, our results challenge our current understanding of the interaction between the microbiome and mosquito development.

## Results

### Testing mosquito diets to support axenic mosquito growth

In order to define the nutritional requirements needed to support mosquito development, we tested multiple mosquito diets. Mosquito diets free of living bacteria generally resulted in widespread mortality or stalled larval development (Table 1). In contrast, the majority of gnotobiotic larvae (colonized by *E. coli* strain K12) survived to adulthood. Yet, supplementing the mosquito diet with heat killed or sonicated *E. coli* cells in the medium failed to rescue development (Table 1). Furthermore, the highest concentrations of heat-killed or sonicated *E. coli* cells resulted in larval mortality. A mixture of amino acids and vitamins commonly used for cell culture were also assessed for their ability to support larval growth. Larvae failed to develop at low concentrations of the added nutrients, whereas the mixture was lethal to the larvae at high concentrations (Table 1). To ascertain if the ability to rescue larval development was unique to bacteria, we inoculated the standard larval diet with 100 μl of an active baker’s yeast culture and found that this too rescued development. However, solutions of 1% or 10% sterilized yeast extract did not rescue development, with 10% yeast extract causing larval mortality. These data suggest that both live bacteria and fungi are both capable of rescuing larval development. Yet a common observation across the different diets was that in high concentrations many of the supplements were toxic to the larvae (Table 1), suggesting mosquito larvae are sensitive to high concentrations of particular compounds in their environment.

**Table 1.**
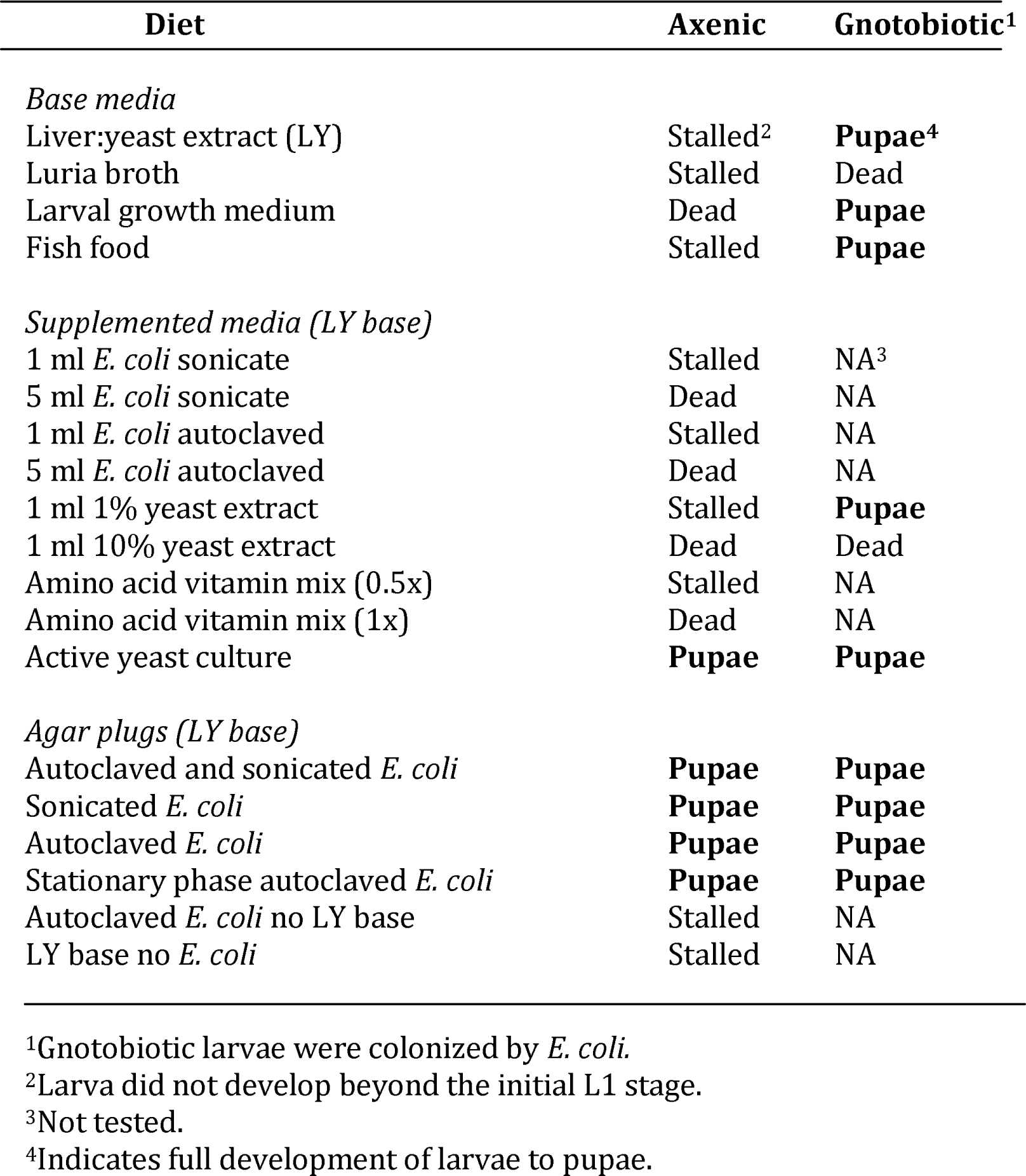
Effect of sterile diets on larval development.

| **Diet** | **Axenic** | **Gnotobiotic<sup>1</sup>** |
| --- | --- | --- |
| *Base media* |  |  |
| Liver:yeast extract (LY) | Stalled<sup>2</sup> | **Pupae<sup>4</sup>** |
| Luria broth | Stalled | Dead |
| Larval growth medium | Dead | **Pupae** |
| Fish food | Stalled | **Pupae** |
| *Supplemented media (LY base)* |  |  |
| 1 ml *E. coli* sonicate | Stalled | NA<sup>3</sup> |
| 5 ml *E. coli* sonicate | Dead | NA |
| 1 ml *E. coli* autoclaved | Stalled | NA |
| 5 ml *E. coli* autoclaved | Dead | NA |
| 1 ml 1% yeast extract | Stalled | **Pupae** |
| 1 ml 10% yeast extract | Dead | Dead |
| Amino acid vitamin mix (0.5x) | Stalled | NA |
| Amino acid vitamin mix (1x) | Dead | NA |
| Active yeast culture | **Pupae** | **Pupae** |
| *Agar plugs (LY base)* |  |  |
| Autoclaved and sonicated *E. coli* | **Pupae** | **Pupae** |
| Sonicated *E. coli* | **Pupae** | **Pupae** |
| Autoclaved *E. coli* | **Pupae** | **Pupae** |
| Stationary phase autoclaved *E. coli* | **Pupae** | **Pupae** |
| Autoclaved *E. coli* no LY base | Stalled | NA |
| LY base no *E. coli* | Stalled | NA |

<sup>1</sup>Gnotobiotic larvae were colonized by *E. coli*.
<sup>2</sup>Larva did not develop beyond the initial L1 stage.
<sup>3</sup>Not tested.
<sup>4</sup>Indicates full development of larvae to pupae.

A notable observation from the above experiments was that while gnotobiotic mosquitoes colonized by *E. coli* were generally capable of development, *E. coli* itself did not grow well in the larval media. Thus, we hypothesized that the majority of the *E. coli* were maintained inside the mosquito midgut rather than in the external environment. This observation, coupled with prior knowledge that axenic *Drosophila* larvae are reared on a solid-state cornmeal agar, which allows for the direct consumption of a highly nutrient-rich diet^37^ led us to develop a mosquito diet with a high concentration of bacterial components embedded in an agar plug. This was the only diet able to support the development of mosquitoes in the absence of live bacteria. Sterility of the resulting larvae and adult mosquitoes was confirmed by both culture-dependent and independent methods (PCR of 16S rRNA genes, see methods) (Figure 1).

**Figure 1:**
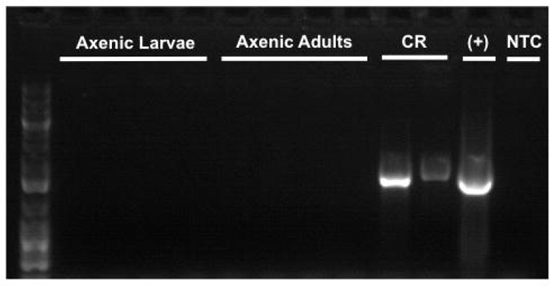
PCR detection of bacterial DNA. Total DNA was extracted from axenic and conventionally reared (CR) mosquitoes and employed as a template for amplification of bacterial 16S rRNA genes.Lanes 1-8 show no visible PCR products from four of each axenic L4 larvae and adult mosquitoes, whereas amplification products were identified in the CR mosquitoes (lanes 9-10). A positive control (+) containing amplified *E. coli* K-12 DNA and a non-template control (NTC) are also included on the gel. Amplification consisted of 30 cycles (see methods).

### Development of axenic, gnotobiotic, and CR mosquitoes

Having generated axenic mosquitoes and confirmed their sterility, we assessed if there were developmental effects associated with the axenic state by comparing axenic mosquitoes with gnotobiotic and conventionally reared mosquitoes (CR). The gnotobiotic group consisted of sterilized larvae colonized by *E. coli* and the CR group was sterilized larvae colonized by bacteria from a conventionally reared mosquito colony (see methods below). Out of the three conditions, the axenic group exhibited the lowest rates of larval mortality (average: 1.85%±1.85%) within the fourteen-day experimental period, while the CR group exhibited the highest rates of mortality (average: 61.1%± 11.1 %). Axenic mosquitoes also exhibited a significant delay in time to pupation when compared to their gnotobiotic counterparts (p<0.0001; Figure 2). On average, axenic larvae pupated approximately three days after the gnotobiotic larvae. There was no significant difference in pupation time between the axenic and CR groups.

**Figure 2:**
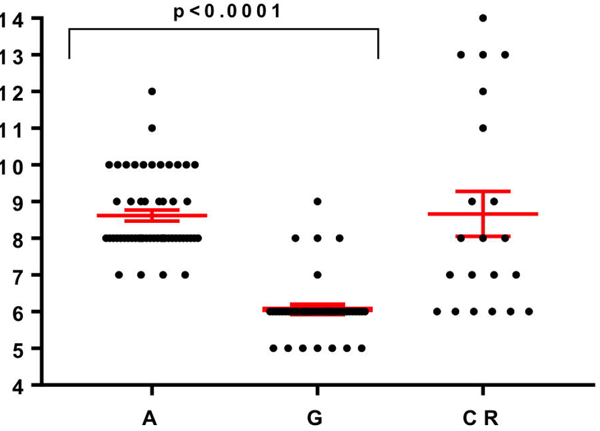
Delayed development in axenic larvae. Time to pupation is shown for axenic (A), gnotobiotic (G), and conventionally reared (CR) larvae. Points represent time to pupation for individual larvae. The red bars signify mean time to pupation for the three replicates and the standard error.

The relatively rapid development of the gnotobiotic group is notable because *E. coli* K-12 is a laboratory-adapted bacterium that carries mutations in lipopolysaccharide and amino acid synthesis, making it a poor colonizer of most hosts^38^. *E. coli* K-12 is therefore unlikely to be a normal member of the mosquito microbiome. Yet, we observed faster larval growth in the presence of a mono-culture of *E. coli* than in the normal microflora of laboratory-reared *Ae. aegypti* (Figure 2). This raises interesting questions regarding the interactions between the diversity and composition of the host’s microbiome and the host’s phenotype. However, the high mortality and longer development times of the CR group in comparison to the gnotobiotic group is likely attributable to the nature of our experimental setup. Larvae were reared individually in six-well plates, providing an ideal environment for growth of the native mosquito microbiota, which in turn caused widespread mortality in larvae in the CR group. Presumably, competition between the mosquitoes and bacteria for resources, or the production of inhibitory compounds by the bacteria were responsible for larval mortality. Additionally, the axenic group was the only group that did not exhibit any mortality during the pupal stage, which further supports that mortality may be related to the bacterial load in the system. Because the rates of mortality were unusually high and did not reflect larval survivorship under normal mosquito rearing conditions, the CR group was excluded from the phenotypic analyses described below.

### Biometric assessment of axenic and gnotobiotic mosquitoes

To determine if our axenic rearing conditions altered mosquito phenotypic traits, we examined adult sex ratio, wing length, and survivorship. The axenic group contained a greater proportion of males than the gnotobiotic group, although the difference was not significant (p=0.23, Figure 3a). These results were somewhat surprising considering that a previous study demonstrated that *Culex molestus* mosquitoes reared on low concentration larval diets had higher proportions of males to females compared to those reared on a high concentration larval diet^39^. Interestingly, they also observed larger males and females, as measured by wing length, in the high versus low concentration larval diets, whereas we observed no significant differences in wing length (mm) among the males and females in the axenic and gnotobiotic groups (unpaired t-test, males: t(61) = 0.668, p=0.506, females: t(37)=0.114, p=0.99; Figure 3b). Finally, there was no observable effect on adult survivorship after 14 days (data not shown).

**Figure 3:**
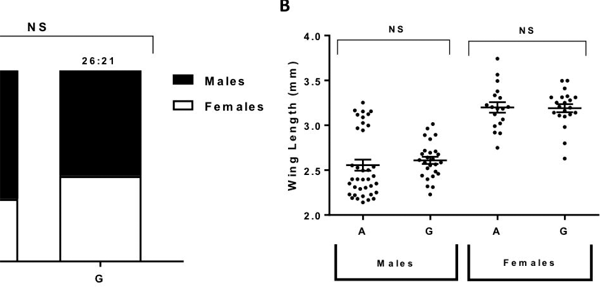
Biometric assessment of adult mosquitoes. (a) Male:female ratio of adults for axenic (A) and gnotobiotic (G) mosquitoes. Using Fisher’s exact test, no significant difference in sex ratio was identified (p>0.05) between the two groups. B) Wing length of adults for axenic and gnotobiotic mosquitoes. Points represent individual mosquitoes. Mean wing length and standard error of all individuals are signified by the black bars. No significant difference in wing length was observed between the two groups (unpaired t-test; p>0.05).

### Egg clutch size of axenic mosquitoes

Conventionally reared females obtained from our laboratory-maintained colony had larger egg clutch sizes (74.29 ± 3.26, n=21) than their axenic counterparts (60.75 ± 3.34, n=16; unpaired t-test, t(35)=2.848, p=0.007; Figure 4), a reduction of approximately 18% fewer eggs. The reduced clutch sizes did not affect the viability of the eggs as larvae laid by axenic females readily hatched. Sterility of the subsequent generation was confirmed by culture-dependent and independent means. These data indicate that under the proper conditions mosquitoes could be reared axenically over multiple generations, leading to the potential establishment of axenic mosquito colonies.

**Figure 4:**
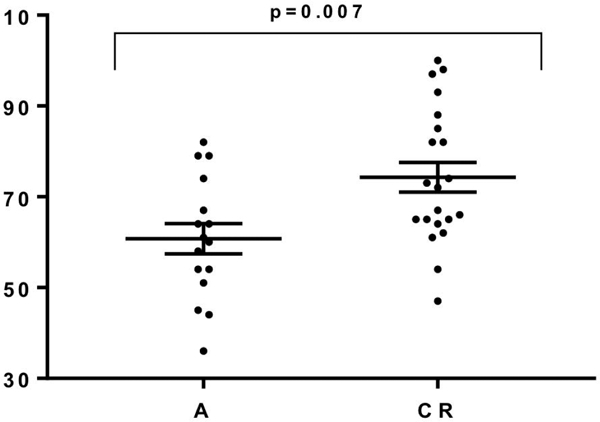
Decreased egg clutch size in axenic mosquitoes. Mean egg clutch size and distribution is shown for axenic (A) and conventionally reared (CR) mosquitoes. Points represent clutch size for individual females. Mean clutch size and standard error of all individuals are signified by the black bars. Clutch size in CR mosquitoes was significantly greater than in axenic mosquitoes (unpaired t-test; p=0.007).

### Characterizing bacterial components that rescue mosquito development

The development of agar plugs containing liver:yeast extract and infused with attenuated *E. coli* was based on the hypothesis that large populations of actively growing bacteria were present in the mosquito midgut, and these populations of bacteria were producing the required nutrients to rescue larval development. Yet, the cellular components that rescued development were unknown. The initial agar plugs used in this study contained a mixture of sonicated, filter sterilized, and autoclaved *E. coli* cells derived from a mid-log phase culture. These agar plugs rescued larval development, and are the basis of the results presented above.

To better define the requisite bacterial components in the agar plugs needed for development, several cell fractions were tested for their ability to support larval development. Both sonicated and filter sterilized *E.coli* cells and autoclaved cells alone rescued larval development (Table 1), indicating that the cellular components supporting development were not heat labile. The initial agar plugs were made from mid-log phase cells based on the theory that the nutrients required for development were obtained from metabolically active and vigorously growing cells. However, agar plugs derived from stationary phase *E. coli* were similarly successful in rescuing development. Agar plugs consisting of the bacterial components but without the liver:yeast extract base were also tested. In this case, the agar plugs failed to rescue development (Table 1). Finally, agar plugs containing the liver:yeast extract base but no attenuated *E. coli* did not support larval development, indicating the effect of the agar plugs was not due to the effects of a solid diet or nutrition provided by the agar.

Taken together, the above observations indicate that the components of the bacterial cells being utilized by the mosquitoes are heat stable and likely include some of the larger cellular components like the cell membrane, these components are present throughout the bacterial cell cycle, and, while required for larval development, they are not sufficient on their own to support full larval growth. Identifying the specific molecules supplied by the bacteria to fuel larval development will be a critical step in designing a fully synthetic defined medium for the rearing of axenic mosquitoes.

## Discussion

Contrary to previous findings^27,35,36^, we demonstrate that it is possible to rear axenic mosquitoes under laboratory conditions. Furthermore, the colonization of axenic larvae to generate gnotobiotic mosquitoes demonstrates an ability to manipulate the microbiome, specifically to imprint axenic mosquitoes with bacterial communities of a known composition. This transitions the microbiome to an experimental variable that can be utilized to gain a better mechanistic understanding of the interaction between the microbiome and the phenotype of the host.

Here we show that bacterial components, when provided in high concentrations in a semi-solid form rather than freely in the mosquitoes’ aquatic environment, can rescue larval development. This suggests that the primary association between mosquitoes and their gut microbiota is nutritional rather than symbiotic. Additionally, this relationship is not unique to bacteria, as baker’s yeast can also rescue development (Table 1). Finally, a companion paper to this study demonstrates that axenic larvae can be rescued through feeding on a diet consisting of cell-cultured live mosquito and *Drosophila* cells (Conor McMeniman, personal communication). In sum, these observations suggest that mosquitoes are unable to produce essential nutrients on their own, and that in nature these nutrients are supplied by the microbiome. A study of the transcriptional differences between axenic and colonized larvae may support this hypothesis. Axenic larvae (unable to develop due to a lack of a microbiome) displayed significant down-regulation of peptidase genes and an upregulation of amino acid transporters in comparison to their microbially colonized cohorts^40^. This suggests that protein and amino acid metabolism is significantly altered in axenic larvae. Yet when we supplemented the larval diet with an amino acid and vitamin mixture, low concentrations were insufficient to rescue larval development and high concentrations were lethal (Table 1). Thus, a diet with the appropriate concentration of amino acids and proteins appears to be critical for larval development, but we have so far been unable to identify the necessary components or concentration for a fully synthetic larval diet.

Previously, bacterial-mediated hypoxia was identified as a potential mechanism to explain the apparent requirement of live bacteria for mosquito development^36^. This report was based on the observation that *E. coli* mutants in cytochrome *bd* oxidase could not rescue larval development^36^. This oxidase has a high affinity for oxygen, and allows facultative anaerobes to maintain aerobic respiration under low oxygen conditions^41^. In this regard, the role of this enzyme complex is generally in response to anaerobic conditions, rather than a cause of them. Thus, the inability of mutants in this complex to rescue larval development may be due to an inability of cytochrome *bd* mutants to survive anoxic conditions. Furthermore, cytochrome *bd* oxidase complexes can protect bacteria from agents synthesized by the host immune system,such as reactive oxygen species and nitric oxides^42,43^. Therefore, cytochrome *bd* oxidase mutants may be compromised in their ability to colonize host organisms^44^. Finally, our data show that disrupting *E. coli* cells, either through sonication or autoclaving and feeding them to mosquitoes, can rescue larval development, suggesting that larval development can occur without bacterially-mediated anoxia. Collectively, these observations support the hypothesis that the ability of the microbiome to rescue larval development is based on a nutritional relationship rather than an active interaction between the bacteria and the host.

The importance of the microbiome in the functional biology and physiology of vertebrates and invertebrates has been well documented^1,2^. It is unsurprising, therefore, that axenic mosquitoes exhibit phenotypic differences when compared to gnotobiotic or CR mosquitoes. Axenic mosquitoes demonstrated a delay in development when compared to the gnotobiotic group. A similar delay in development has been observed in other axenic organisms, including *Caenorhabditis elegans* and *Drosphila*^45^’^46^. In the case of *C. elegans,* an increase in development time is usually coupled with an increase in longevity, with axenic organisms living longer than organisms reared on a non-axenic culture^45,47^. This pattern is not observed in axenic *Drosophila,* the longevity of which appears to be diet-specific^48,49^. This suggests that the removal of the microbiome has differing effects on host biology and demonstrates why expanding the number of axenic models will help to uncover the common and diverging effects of the microbiome on host physiology. It is interesting to note that no significant difference in wing length was observed between axenic and gnotobiotic mosquitoes, suggesting that axenic mosquitoes are able to compensate for an increase in developmental time and reach full body size once a certain threshold in development has been surpassed.

Axenic mosquitoes also displayed a significant decrease in fecundity. A study on *Aedes aegypti* mosquitoes that were colonized by single bacterial strains showed that differences in fecundity of gnotobiotic females can be strain-specific^50^. Our results suggest that an absence of bacteria during development can detrimentally affect female egg clutch size. It is important, however, to note that mosquito egg clutch size has also been linked to nutrition^51,52^, which could explain the observed differences between the axenic and colonized groups. Bacterial turnover may supply the colonized mosquitoes with a steady source of nutrients, which is not available to the axenic group. Additionally, the microbiome is associated with an increased ability to extract energy from food^53^. Therefore, we would expect that this would lead to a better nutritional status for gnotobiotic and CR mosquitoes, possibly explaining reduced egg clutch size for the axenic mosquitoes.

In summary, this study presents a method to rear axenic *Aedes aegypti* mosquitoes from eggs to adults and into the subsequent generation in the complete absence of a microbiome. We show that axenic mosquitoes develop normally, but with a delay in the time of development. Axenic mosquitoes show decreased mortality and smaller egg clutch sizes in comparison to their bacteria colonized cohorts. As mosquitoes are a major global health concern, interventions that could decrease mosquito fecundity are a common objective for mosquito management. The data presented here suggest that the microbiome may be a potential target for future control strategies. Using bacteria as a tool in mosquito control, a method referred to as paratransgenesis^15,56^, has already been pursued. However, these studies have thus far been hampered by a lack of effective tools to manipulate the microbiome. The methods presented in this study add mosquitoes to the collection of organisms for which an axenic state can be maintained, underpinning our ability to treat the microbiome as a controlled experimental variable in organismal studies.

## Methods

### Preparation of Mosquito Rearing Substrates

Multiple diets and supplements were tested for their ability to support the development of axenic mosquitoes. The first group of treatments was based on the standard diet for the colony raised mosquitoes, which consisted of a 0.1% solution of 3 parts liver extract (Difco, dessicated, powdered beef liver) and 2 parts yeast extract (Fisher Scientific, granulated yeast extract). The standard diet was also supplemented with the following: 5 ml Luria Broth (LB), 1 ml or 5 ml of an overnight culture of sonicated *E. coli* cells (throughout the manuscript *E. coli* refers to the wild-type strain K-12^54^), 1 ml or 5 ml autoclaved *E. coli* cells (overnight culture), 0.2% or 2% (w/v) yeast extract, 100 μl of an overnight culture of live baker’s yeast *(Saccharomyces cerevisiae),* and a 1x and 0.5x solution of an amino acid (Gibco MEM Amino Acids 50x stock) and vitamin (Gibco Vitamin Solution 100x stock) solution mixture. Two diets included a media base not consisting of the standard diet. These were: 0.1% sterile fish food (TetraminTropical Flakes) and a synthetic larval growth media previously described^33^.

The final diet tested was a mixture of sonicated and heat-killed *E. coli* embedded in agar plugs. A starter culture of *E. coli* was grown overnight at 37°C and used to inoculate two flasks of 500 ml of LB broth. To harvest mid log-phase cells, the inoculated flasks were placed on a shaker and incubated at 37°C for approximately 5 hours to an OD of ~0.8. The cells were then centrifuged at 1100 rpm for 10 minutes. After discarding the supernatant, the bacterial pellet was re-suspended in 20 ml of PBS and sonicated at 65% amplitude using a Fisher Scientific Model 120 Dismembrator for a total of 3 minutes. The sonicated cells were then centrifuged at 4000 rpm for 10 minutes. The supernatant was filter sterilized using a 0.2 μm PES membrane filter, and the pellet was re-suspended in 10 ml of sterile water and subsequently autoclaved at 121°C for 30 minutes. The resultant filtrate and autoclaved pellet were then combined with 30 ml of a 1.5% agar solution containing 3.3% 3:2 liver:yeast extract. The agar mixture was poured into standard Petri dishes and stored at 4°C.

For the plugs that contained only sonicated and filter sterilized *E. coli,* the above standard protocol for culturing and sonicating the bacteria was followed; however, 10 ml of sterile water was substituted for the autoclaved bacterial pellet. Similarly, for plugs that contained only mid-log phase autoclaved *E. coli,* the standard protocol was followed with the following exceptions: after re-suspension in 20 ml of PBS, the pellet was autoclaved at 121°C for 30 minutes and was then combined with 40 ml of a 1.5% agar solution containing 3.3% 3:2 liver:yeast. For plugs containing stationary phase *E. coli,* two 500 ml flasks of LB were inoculated with *E. coli* and allowed to grow on a shaker at 37°C overnight. The culture was then centrifuged, re-suspended, autoclaved, and combined with agar and liver:yeast extract as detailed above. A full procedure documenting the making of agar plugs is attached as a supplemental file to this manuscript.

### Egg Sterilization

*Aedes aegypti* eggs were acquired from a colony of laboratory-reared mosquitoes (Orlando strain) maintained in environmental chambers at 28°C with a 16:8 light:dark photoperiod. Sterilization of the *Aedes aegypti* eggs was carried out as previously described.^35^ Briefly, a small segment of egg-covered filter paper was washed for five minutes in 70% ethanol, followed by a five-minute wash in a 3% bleach and 0.2% ROCCAL-D (Pfizer) solution, and an additional five minute wash in 70% ethanol. The sterilized eggs were then rinsed three times in autoclaved distilled water and placed in Petri dishes filled with phosphate buffered saline (PBS). Eggs were hatched in a vacuum oven (Precision Scientific Model 29) at 25Hz for 15 minutes at room temperature. A schematic diagram showing the sterilization procedure is shown in Figure 5.

**Figure 5:**
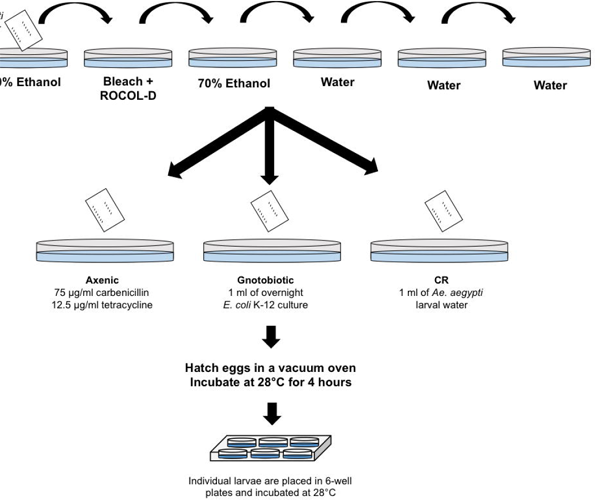
Schematic diagram of egg sterilization. *Ae. aegypti* eggs were collected from colony reared mosquitoes and surface sterilized as depicted. (a) Eggs were serially washed for 5 minutes in each solution. (b) Surface sterilized eggs were transferred to a Petri dish containing sterile PBS and antibiotics for axenic mosquitoes, *E. coli* cells for gnotobiotic mosquitoes, or water collected from colony rearing pans for CR mosquitoes. (c) After hatching the eggs in a vacuum oven and four hours of incubation, individual larvae were transferred from the Petri dishes to individual wells of a six well plate for development assays.

### Bacterial colonization of sterile larvae

The sterile hatched eggs were split into three different treatment groups. Axenic larvae were incubated at 28°C in the presence of 75 μg/ml carbenicillin and 12.5 μg/ml tetracycline for 4 hours. Gnotobiotic larvae were exposed to a 1 ml aliquot of an overnight culture of *E. coli* for 4 hours at 28°C. Finally, the conventionally reared (“CR”) group were inoculated with a 1 ml aliquot obtained from pans of water used to rear larvae from a laboratory maintained *Ae. aegypti* colony. In this manner, each group went through the sterilization procedure, ensuring that any observed differences in mosquito development were not due to effects from surface sterilization of the eggs.

### Rearing of larvae

For each tested condition, six individual larvae were transferred from the Petri dishes to individual wells of a 6-well plate. Each well of the plate contained 5 ml of the rearing substrate, or, in the case of the agar plugs, a 0.4 g plug in a 5 ml solution of sterile saline. Development, time to pupation, and mortality were recorded each day for 14 days after hatching for a total of three replicate plates (i.e. 18 individuals).

### Confirming the Sterility of Axenic Mosquitoes

A subset of the adults that were reared under axenic conditions was tested for the presence of live bacterial cell or bacterial genomic DNA via culturing and 16S rRNA gene PCR. Individual mosquitoes were transferred to a round bottom tube containing a steel BB and 150 μl of sterile PBS and homogenized for 30 seconds at 30 1/s using a Mixer Mill. 50 μl of each homogenate was inoculated into a 14 ml culture flask containing 2 ml of LB broth and incubated 48 hours at 28°C. Negative results were confirmed by an absence of bacterial growth.

For the remainder of the homogenate, total DNA was extracted using a PowerSoil DNA Isolation kit (MoBio Laboratories, Inc.). DNA extractions were PCR amplified using standard 16S rRNA gene primers (27F and 1492R^55^), using the following protocol: initial denaturation at 95°C for three minutes, followed by 30 cycles consisting of 95°C for 45 seconds, annealing at 55°C for 45 seconds, and extension at 72°C for 1 minute 45 seconds, with a final extension at 72°C for 10 minutes. PCR results were verified by agarose gel electrophoresis. Although readily detected in colonized mosquitoes, bacterial DNA amplification was not detected by gel electrophoresis in axenic larvae and adult mosquitoes (Figure 1), despite *E. coli* DNA likely being present in the initial agar plugs. These data were collected from L4 larvae, thus incubation at 28°C, coupled with the presence of live mosquito larvae, presumably broke down any *E. coli* DNA present in the agar plugs. PCR of initially sterilized larvae and the growth media components were also consistently negative by bacterial 16S rRNA gene PCR.

To ensure that the axenic larvae were truly bacteria free we employed a three step verification of sterility: 1) In each experiment, a “sterile” control group of larvae was maintained. This group was processed in parallel to the other treatment groups and fed on the same batch of liver:yeast extract. Any larval development in this group past the L1 stage was taken as an indication of contamination and the experiment was discarded; 2) In each experiment, a subset of axenic larvae and adult mosquitoes were tested for contamination by culturing. A positive test for bacterial presence in any experiment indicated contamination and the experiment was discarded; 3) A subset of axenic larvae (L4 growth stage) and newly emerged adult mosquitoes were tested for bacterial DNA through PCR of bacterial 16S rRNA genes. A positive test for bacterial presence in any experiment indicated contamination and the experiment was discarded.

### Mass Rearing and Blood Feeding Axenic Mosquitoes

After eggs were sterilized and hatched, larvae were incubated for four hours at 28°C in PBS supplemented with 12.5 μg/ml tetracycline and 75 μg/ml carbenicillin. Approximately 150 larvae were placed in a sterile 500 ml polypropylene Nalgene jar containing 125 ml of distilled water and 4 grams of the *E. coli* agar food. After pupation, larvae were allowed to emerge into a UV-sterilized mosquito emergence chamber. Adult mosquitoes were fed filter sterilized 10% sucrose, and female axenic mosquitoes were blood fed sterile defibrinated sheep blood using a Hemotek membrane feeder in a biosafety cabinet. Hemotek feeders were autoclaved and UV-sterilized parafilm was used for the feeds. Similarly, all glassware and forceps used to sort mosquitoes were autoclaved. CR mosquitoes were acquired from the laboratory colony and blood fed using a circulating bath membrane feeder.

### Biometric Assessment of Axenic Mosquitoes

Using the method of mass rearing described above, an equal number of sterile and nonsterile larvae were placed in separate polypropylene jars, allowed to emerge, and were blood fed using a Hemotek membrane feeder. Axenic larvae were fed 10 ml of the *E. coli* agar food and the CR mosquitoes were fed a 1% 3:2 liver: yeast extract solution. After blood feeding, axenic (n=16) and CR (n=21) females were individually placed in autoclaved 50 ml tubes containing sterilized egg-laying filter paper and water. After oviposition, the filter papers were removed from the tubes and the eggs counted.

To assess differences in wing length between the three groups axenic, gnotobiotic, and CR larvae were reared in 6 well plates as per the development assay. After emergence, males and females were knocked down on ice and wings were removed using forceps. Wings were then visualized using a Zeiss Axioplan 2 universal microscope and wing length was measured using Axiovision (v.4.8.1) software.

## Acknowledgements

We thank John Shepard and Michael Thomas for assistance with mosquito colony maintenance. This work was supported in part by grants from the Centers for Disease Control and Prevention (U50/CCU116806-01 and U01/CK000509-01), the US Department of Agriculture Hatch Funds and Multistate Research Project (CONH00773 and NE1443) and the National Institute of Health, National Institute of Allergy and Infectious Diseases (5K22AI099042-02).

